# Wearable Eye-tracking for Research: Automated dynamic gaze mapping and accuracy/precision comparisons across devices

**DOI:** 10.1101/299925

**Authors:** Jeff J. Macinnes, Shariq Iqbal, John Pearson, Elizabeth N. Johnson

## Abstract

Wearable eye-trackers offer exciting advantages over screen-based systems, but their use in research settings has been hindered by significant analytic challenges as well as a lack of published performance measures among competing devices on the market. In this article, we address both of these limitations. We describe (and make freely available) an automated analysis pipeline for mapping gaze data from an egocentric coordinate system (i.e. the wearable eye-tracker) to a fixed reference coordinate system (i.e. a target stimulus in the environment). This pipeline allows researchers to study aggregate viewing behavior on a 2D planar target stimulus without restricting the mobility of participants. We also designed a task to directly compare calibration accuracy and precision across 3 popular models of wearable eye-trackers: Pupil Labs 120Hz Binocular glasses, SMI ETG 2 glasses, and the Tobii Pro Glasses 2. Our task encompassed multiple viewing conditions selected to approximate distances and gaze angles typical for short- to mid-range viewing experiments. This work will promote and facilitate the use of wearable eye-trackers for research in naturalistic viewing experiments.

## Introduction

Eye-tracking offers insights into otherwise hidden aspects of visual processing and cognition. Traditionally, methods for recording eye-movements and gaze behavior have relied on search coils (embedded in contact lenses or implanted surgically) or infrared systems like Dual Purkinje Tracking and screen based eye-trackers (Collewijn and Kowler 2008; Rucci et al. 2007; Epelboim et al. 1995). These techniques offer high-precision and accuracy, but sacrifice ecological validity under naturalistic viewing conditions. Screen based eye-trackers, for instance, offer an ease-of-use that has made them a popular choice for psychology labs, but severely limit the nature and quality of experimental stimuli; stimuli are limited to two-dimensional digital representations, and must be sized to fit within the screen dimensions. Moreover, subjects are often seated at a fixed distance from the screen with limited head and body mobility, which can be a difficult position to sustain for extended periods of time (and in particular for studies involving children or toddlers). These methods remain powerful approaches to vision research, providing high-precision and accuracy, but preclude studying gaze behavior in more naturalistic viewing contexts.

Recently, wearable eye-trackers have emerged as an alternative approach, allowing researchers to record gaze and pupillometry measures as subjects freely navigate an environment. With wearable trackers, the pupil cameras are attached to the frame of a pair of eyeglasses that subjects wear while they move about untethered. As such, wearable devices hold the exciting potential of bringing eye-tracking to new contexts and new subject populations. Michael Land built one of the first portable and wearable eye trackers almost 25 years ago, but its use was limited because eye position had to be calculated offline due to computational limitations at that time (M. F. Land and Lee 1994; M. F. Land and Furneaux 1997; M. Land, Mennie, and Rusted 1999). The use of wearable eye-tracking devices for research continues to be hindered by the significant analytic challenges that are introduced, as well as by a lack of published benchmarking routines, which can be used to compare performance measures among the competing devices on the market.

Here, we address these shortcomings. This article has two independent objectives: 1) introduce automated analysis methods that facilitate the use of wearable eye-trackers in research contexts by mapping gaze locations to planar targets in the environment, and 2) use these methods as a means to compare calibration accuracy and precision between 3 popular models of wearable eye-trackers.

The first objective is achieved by leveraging advances in computer vision algorithms that have made it possible to automate many of the analytic challenges that wearable devices introduce. We outline those challenges below and describe specific steps taken to resolve them. Our approach relies entirely on freely available, open-source software, and is well-suited for gaze mapping to static 2D physical stimuli in the environment. Furthermore, we make available the specific processing pipeline we have developed as part of our efforts.

Our second objective is to benchmark calibration accuracy and precision across 3 popular models of wearable eye-tracker. In order for wearable eye-trackers to be practical for research studies, it is critical to know how well mapped gaze locations correspond to where a subject was actually looking. We designed a task to measure the accuracy and precision with which a measured gaze location can be mapped to a known point in the environment. We explicitly designed this task to approximate conditions under which wearable eye-trackers could be used for indoor, short-to-medium viewing range, research contexts. Namely, subjects were presented with a physical target stimulus affixed to a wall, and measured performance at distances ranging from 1-3*m*, and gaze angles spanning +/-10° relative to a point centered in the field of view. We report our findings below with additional practical details for using wearable eye-trackers.

## Automated Dynamic Gaze Mapping

### Overview and challenges

The tools we present solve a major issue in using wearable devices in a research context: translating a recorded gaze location from an egocentric reference frame to a fixed reference frame (e.g. a fixed planar stimulus in the environment). With traditional screen-based eye-tracking, the cameras that record pupil location/size are fixed relative to the screen on which stimuli are presented. Thus, translating gaze locations to the screen’s coordinate system only requires a single linear transformation that maps the camera location to the screen. The appropriate transformation is calculated during calibration steps that typically precede data collection. After calibration, that single transformation is then easily applied to all subsequent data, resulting in gaze locations expressed as pixel locations in terms of screen coordinates, simplifying stimulus-based analyses.

In contrast, wearable eye-trackers record gaze locations relative to the subject (or, more specifically, relative to an outward facing camera affixed to the glasses, which records the subject’s point of view; henceforth *world camera).* In this case, the purpose of the calibration is to produce a transformation that maps recorded gaze locations to an egocentric coordinate space defined within the world camera video frame. Consequently, gaze locations are expressed as pixel locations in a coordinate system that lacks reference to any external features of the environment.

While this configuration allows a subject to freely navigate around an environment, it can pose significant challenges for any analysis that attempts to summarize aggregate viewing behavior on a fixed object in that environment. Particular features of an object will appear in different positions in the world camera as subjects assume different orientations and locations. For example, if a subject were asked to view the Mona Lisa, the perceived location of her lips would depend on the subject’s vantage point, and would constantly shift, relative to the world camera frame, as he or she moved.

In order to study object-based viewing behavior, it is necessary to know *when* a subject fixates on the object, and *where* on the object they are looking. For wearable eye-trackers, this means going through the world camera recording, frame by frame, manually identifying where in the frame the object appears (if it appears at all), and determining whether (and if so, where) the corresponding recorded gaze location(s) fall on the object. Attempting to complete this type of analysis manually is at best tedious, and at worst imprecise and non-reproducible. Existing attempts to automate this procedure have relied on placing detectable landmarks (e.g. fiducial markers) in the environment (Walker et al. 2017). This approach can expedite analysis, but may be introducing elements to the environment which bias subjects’ viewing behavior. A better approach would be to detect salient features of the target stimulus itself to use as reference landmarks on each frame of video.

This approach can be achieved and automated by adapting modern computer vision algorithms designed to detect corresponding features between two images. Feature matching takes pairs of images and begins by identifying independent sets of local keypoints on each. Depending on the particular algorithm, keypoints may be defined on the basis of contrast, neighboring pixel values, or image topology. The two sets of keypoints from each source image are compared to determine which, if any, occur in both images. With non-identical source images, the keypoint descriptors will obviously vary between images. The criteria for determining whether keypoints “match” can be tuned by the user, and may vary according to the particular algorithm used, but in general are based on the Euclidean distance between descriptors (see ((Lowe 2004)) for a more in-depth discussion). The result is a list of matching keypoints, where each contains two pairs of coordinates: one for a point’s pixel location in the original image, and one for the pixel location in the comparison image.

The next step is to use the corresponding key point coordinates to produce a transformation that maps between the coordinate systems of the two source images. Any two images that depict the same 2D surface (for instance, two photographs of the Mona Lisa, taken from different angles) can be aligned by means of a rigid body transformation comprised of scalings, translations, rotations, and perspective shifts. The set of corresponding coordinates produced by feature matching can be used to solve for the unique rigid body transformation that will align the two original images. That transformation is encoded as a linear transformation and represented as a matrix that can be used to map between the coordinate systems of the two original images. To minimize errors introduced by imprecise keypoint matching, the transformation calculation can include methods to estimate inlier vs. outlier keypoints (i.e. random sample consensus; RANSAC).

### Feature Matching with Wearable Eye-Trackers

Feature matching algorithms can be adapted to wearable eye-trackers in order to produce a robust and efficient automated analysis pipeline. In this case, feature matching will be used to find corresponding keypoints between a high-resolution reference image and a single frame extracted from the world camera video. Once a transformation has been obtained between the frame and reference image, the gaze data on that frame can be mapped onto the reference image. Thus, applying this approach to an entire video recording enables the gaze locations to be dynamically mapped on a frame-by-frame basis to a fixed reference image.

The reference image should be a digital version of the target stimulus that subjects will be asked to view during data collection, tightly cropped to the borders of the stimulus. The reference image keypoints (henceforth *reference keypoints)* need only be obtained once at the beginning of the analysis. Once eye-tracking data has been collected, feature matching proceeds independently on each frame of the world camera video. On each frame of the video, the steps are:

1. 1. Identify matching keypoints (if any) between video frame and reference image.
2. 2. Obtain a linear transformation matrix mapping from the world-camera frame coordinates to the reference stimulus coordinates
3. 3. Apply the linear transform to that frame’s gaze data to map gaze position onto the reference image.

We developed software tools to automate each of these steps using the popular open source computer vision library OpenCV (Bradski and Kaehler 2008). Each of these steps is described in greater detail below.

*Code for our automated gaze mapping pipeline can be found online at: https://github.com/ieffmacinnes/glassesCalibration/tree/master/gazeMappingPipeline*

*For tutorials and reference on feature-matching in OpenCV, see:*

https://docs.opencv.org/3.0-beta/doc/pytutorials/pyfeature2d/py table of contents feature2d/py t able of contents feature2d.html

#### Identify matching keypoints

The first step is to find a set of keypoints for each of the two images that you wish to map between. In an eye-tracking context, one of those images will always be the high-resolution reference image; thus, once the reference keypoints are identified, they can be re-used repeatedly throughout the analysis. The comparison image will be a single frame extracted from the world camera video, and will require a unique set of features for every new frame.

The particular keypoints for each image are obtained using the Scale Invariant Feature Transform algorithm (SIFT) (Lowe 2004). We chose the SIFT algorithm due to its resilient to the types of image transformations that this analysis presents; namely, rotations, scaling, and perspective shifts. In addition, SIFT creates a rich description for each keypoint along 128 dimensions, which helps ensure accurate matches when comparing corresponding keypoints between two images. It should be noted that while SIFT is free for research purposes, any commercial application of the algorithm requires purchasing a license agreement. License-free alternative algorithms exist (e.g. BRISK (Leutenegger, Chli, and Siegwart 2011), FAST (Rosten and Drummond 2006; Rosten, Porter, and Drummond 2010), ORB (Rublee, Rabaud, and Konolige 2011)), and are available within OpenCV, but are not discussed in detail here.

Once keypoints have been identified on the reference and frame images, the next step is to determine which (if any) of the keypoints are shared in common between the two images (**Fig 1.B**). The camera frame keypoints are compared to the reference keypoints using the Fast Approximate Nearest Neighbor search algorithm (FLANN) (Muja and Lowe 2009). The FLANN algorithm can be tuned using a variety of parameters (in our pipeline, we specified 5 trees and 10 iterative searches through the trees; threshold for “match” was set using a Euclidean distance ratio of 0.5). All of the pairs of keypoints that meet criteria for a match are passed along to the next stage of processing.

**Figure 1:**
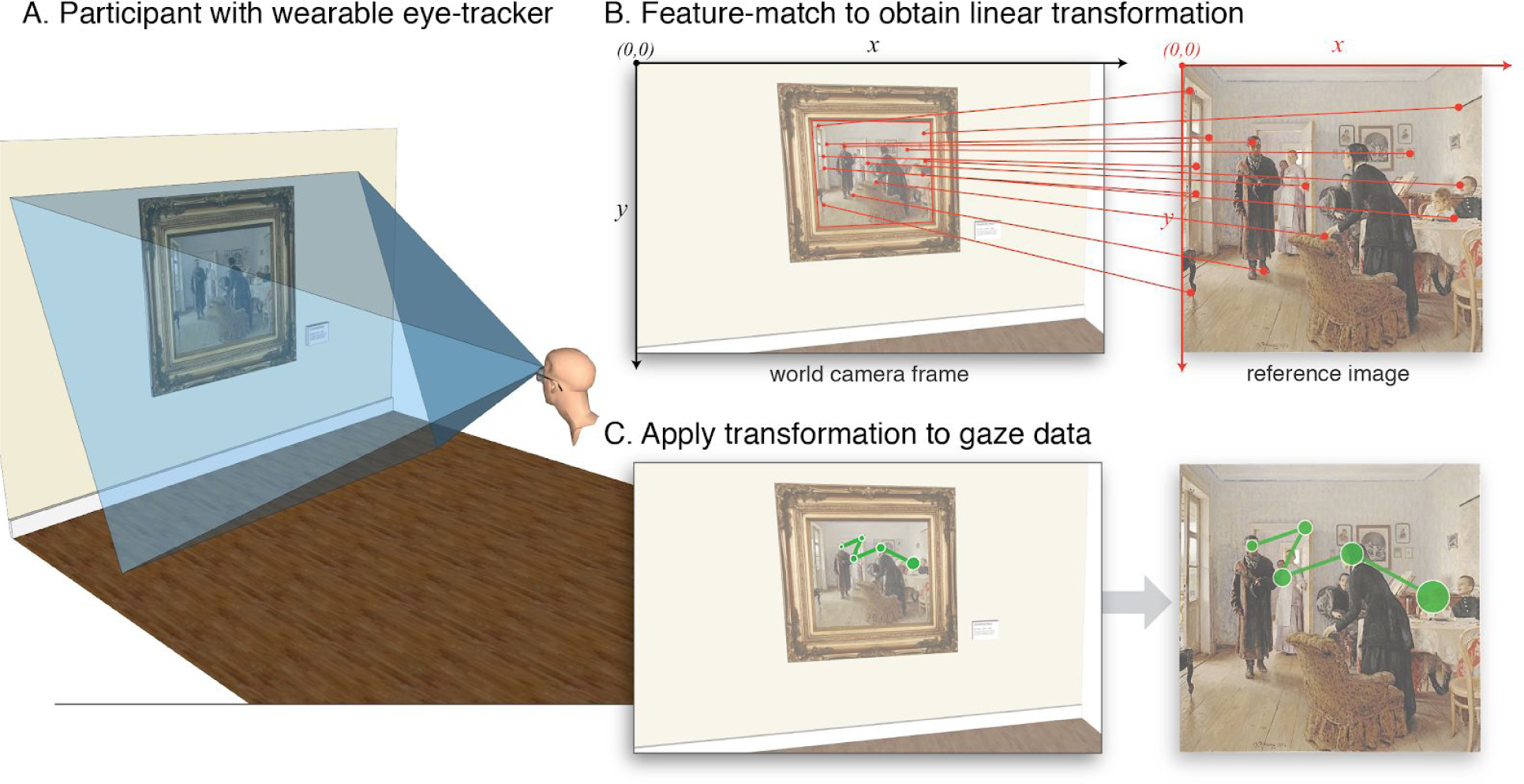
Overview of mapping dynamic gaze data to fixed reference image. **A.** Wearable eye-trackers record data relative to an outward facing world camera capturing a participant’s point-of-view. The location of the target stimulus (in this case, the framed painting) depends on the participant’s position and orientation at each point in time. **B.**. Mapping gaze data to a fixed reference image involves detecting where the target stimulus appears in each frame of the world camera video. Our approach uses SIFT based feature-matching to find corresponding keypoints on both the world camera frame and the target reference image. The set of matching keypoints can be used to obtain a unique linear transformation that maps from a single world camera frame to the reference image. **C.**. The linear transformation for that frame can then be applied to all of the gaze position data corresponding to that same frame. This step yields gaze data expressed in terms of reference image coordinates.

#### Obtain linear transformation

As long as there is a minimum number of matching keypoints (see below), a 2D linear transformation can be calculated that maps from one set of keypoints to the other; that is, from the world camera coordinate system to the reference image coordinate system. In general terms, the relationship between two views of the same 2D planar surface is referred to as its homography, and can be expressed as a linear transformation matrix. The transformation matrix specifies the instructions for mapping from one view to the other via a sequence of 3D rotations and translations. In this case, we use the matching keypoints to find a homography that will transform the keypoints on the camera frame to the corresponding keypoints on the reference image (**Fig 1.B**). Calculating a homography requires only 4 matching keypoints; to identify the keypoints that yield the best homography model, we used RANSAC based outlier detection on the full set of matching keypoints to minimize the influence of imprecise matches.

#### Apply linear transformation to gaze data

Once the homography has been obtained, that same transformation can be used to project new coordinates from the camera frame to the reference image. Since the eye-tracker records gaze data in world camera coordinates, we can use the camera-to-reference image transformation matrix to project the gaze coordinates (corresponding to this frame only) to the reference image coordinate system. The result will be gaze data that is expressed pixel coordinates relative to the reference image, facilitating further analyses (**Fig 1.C**).

Applying this same approach to an entire video recording enables the gaze locations to be dynamically mapped on a frame-by-frame basis to a fixed reference image.

## Comparing Performance Across Devices

A number of manufacturers have released wearable eye-tracking models and marketed them for research purposes. Choosing an appropriate model to fit specific experimental needs is not trivial, as the devices vary in terms of resolution and sampling rates, how they perform calibrations, how they record and format data, and the degree to which they offer low-level access to researchers. Moreover, they can range in price from hundreds to tens of thousands of dollars. Given the range of options available, our next objective was to benchmark the accuracy and precision performance of 3 popular models of wearable eye-trackers on our calibration task, as well as provide a reproducible set of procedures that can be used to benchmark new models in the future.

### Wearable Eye-tracker Models

We selected the Tobii Pro Glasses 2 (https://www.tobiipro.com), SMI ETG 2.6 (https://www.smivision.com), and Pupil Labs 120Hz Binocular (https://pupil-labs.com) wearable eye-trackers and directly compared how well each performed on our calibration task. (Note that SMI was acquired by Apple in June 2017, and will not be releasing new models (Matney 2017)). In addition, since the time of data collection, mobile eye-trackers with faster gaze sampling rates are available from both Pupil Labs (200Hz) and Tobii (100Hz).

#### Calibration Procedures

Each eye-tracker came with its own set of recommended calibration procedures. The Tobii Glasses 2 offer a single calibration method; Pupil Labs and SMI each provide 2 or more calibration methods to suit different experimental configurations. For these models, we chose the method recommended to provide the optimum calibration based on our task design. The specific calibration procedures we followed for each model are described below:

- **Pupil Labs - 120Hz Binocular:** Pupil Labs (Kassner, Patera, and Bulling 2014) offered the most calibration options out of the models tested, including methods tailored to close-range (e.g. computer screen), mid-range, and far-range viewing conditions. We used a “manual marker” based approach, in which participants were instructed to fixate on a handheld target marker as it was placed in various locations throughout their field of view. The markers were held at the same distance from the participant as the task stimuli on that trial (i.e. 1*m,* 2*m,* or 3*m*). Approximately 9 unique locations were used for each calibration, distributed throughout the participant’s field of view. Following calibration, participants were asked to again fixate on the marker as it was moved throughout the scene. Quantified performance measures were not available, and thus in cases of ambiguous-to-poor correspondence between the calibrated gaze point and the marker location (as determined by the researcher), the calibration procedures were repeated until the correspondence improved.
- **SMI - ETG 2.6 glasses:** SMI utilizes natural feature calibration, whereby the participant fixates on a specific location in the environment, which the experimenter then identifies on the live-stream within the software by clicking on the corresponding feature in the world camera image. SMI provides the choice between 1-pt and 3-pt calibration. We used 3-pt calibration to ensure the best calibration accuracy possible. Participants were asked to fixate on a set of 3 standardized locations in the environment. The locations were evenly spaced around a circle centered within a participant’s field of view and with a diameter of 10° of visual angle. The 3 points were located at 30°, 150°, and 270° around the circle (0° at (1,0) on a unit circle). As the circle radius was based on visual angle, the physical location of the points in the room varied according to the distance conditions. Following calibration, participants were asked to again fixate on the same calibration points. Quantified performance measures were not available, and thus in cases of ambiguous-to-poor correspondence between the calibrated gaze point and the marker location (as determined by the researcher), the calibration procedures were repeated until the correspondence improved.
- **Tobii Pro - Glasses 2:** Tobii provided a single calibration method: 1-pt, manual marker calibration. Participants are asked to fixate on a supplied black and white bullseye calibration target held at distance of 0.8-1.2*m*. Tobii software identifies the target in the world camera video automatically. A message appears indicating whether the calibration was successful or not. Following calibration, participants were asked to again fixate on the marker. Quantified performance measures were not available, and thus in cases of ambiguous-to-poor correspondence between the calibrated gaze point and the marker location (as determined by the researcher), the calibration procedures were repeated until the correspondence improved.

## Methods

*Code for the task and entire data analysis pipeline can be found online at: https://github.com/jeifmacinnes/alassesCalibration*

Using the recommended calibration procedures from each manufacturer, we determined how well gaze location could be mapped to a known position in the environment. We tested each eye-tracker across a parameter space that was chosen to approximate real-life, indoor viewing conditions that these devices could be used for in an experimental context. Specifically, we measured calibration performance on a target stimulus positioned at 3 distances (1*m*, 2*m*, and 3*m*), and 3 gaze angles at each distance (target grids centered at -10°, 0°, and +10° horizontal visual angle relative to center of participant’s field of view). Thus, in total, each eye-tracker was tested across 9 conditions spanning +/-10° of gaze angle at distances ranging from 1-3*m* away.

### Task & Stimuli

Each participant (*N*=3) was tested across 27 conditions (3 eye-tracker models, 3 distances, 3 gaze angle offsets). With each condition, participants were seated and asked to remain still for the duration of the task. Participants were positioned at 1, 2, and 3 meters away from the wall. At each distance, a target grid was positioned on the wall such that the center of the grid fell at gaze angles of -10°, 0°, and +10° of horizontal visual angle relative to the participant (**Fig 2**). For each condition, the task and procedures were the same.

**Figure 2:**
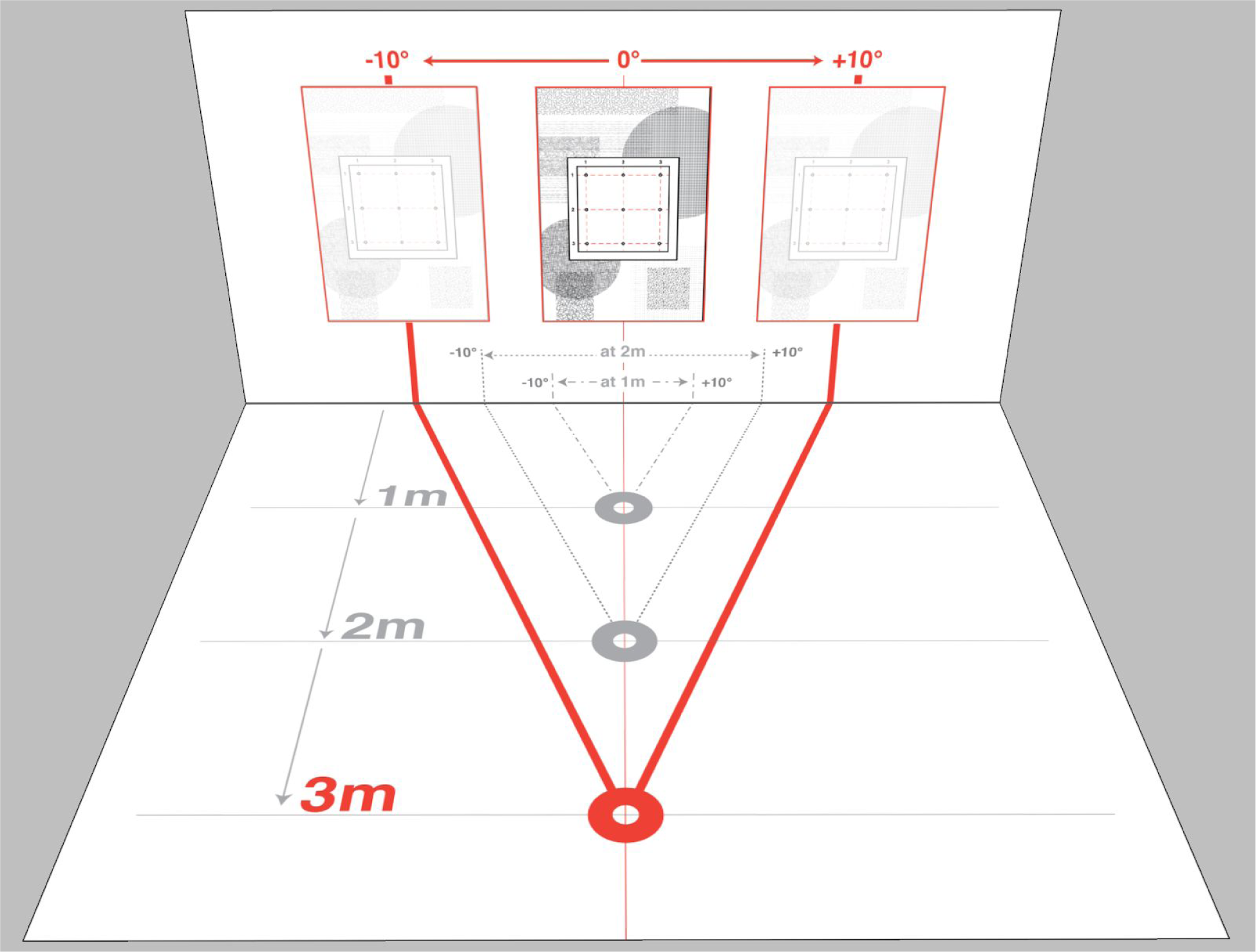
Overview of task conditions. Each participant performed the same task using each of the 3 eye-trackers. The task consisted of 9 unique conditions, spanning a range of distances and gaze angles chosen to approximate a short- to mid-range viewing experiment. At each distance (1*m*, 2*m*, or 3*m*), participants were tested on a target grid positioned at 3 different gaze angles (-10°, 0°, or +10°) relative to the center of their field of view.

Each condition began by calibrating the eye-tracker to the participant according to the manufacturer-specific methods (see *Calibration Procedures* above). The same task was repeated for each condition. Gaze data and camera video recordings were made using the manufacturer specific software and recording devices; a separate dedicated task computer provided visual and audio cues to participants. The task began with the presentation of a start cue that was used to synchronize timing between the recorded gaze data and the trial onsets (see *Time syncing gaze data and world-camera video frames* below). The task consisted of 9 trials in which participants were asked to fixate on a unique location on a target grid. The order and timing of each trial was controlled by a laptop running a customized experimental task (Shinners 2011). Each trial began with the coordinates (column and row number) of the grid location announced via text-to-speech auditory instruction. Participants fixated on each location for 3000 milliseconds. The 9 locations were randomly ordered per condition. It should be noted that the same physical target grid was used in all conditions, and thus the angular subtense between points on the grid varied by distance condition. Our analyses focused on the gaze performance as a function of distance and offset condition, and not on differences between each of the 9 locations on any one condition. As such, our results are labeled according to the gaze offset and distance of the *center* of the target grid at each particular condition.

### Preprocessing

Each manufacturer uses a distinct output format for naming, timestamping, and formatting the output data from its device. We designed manufacturer-specific preprocessing pipelines in order to bring all video and gaze data into a common format to facilitate subsequent analyses. The output from this stage included the world-camera video, a file containing the timestamps of each frame in the world-camera video, and the eye-tracking datafile. Each sample in the eye-tracking datafile contained the timestamp, corresponding world-camera video frame number, and the X,Y coordinates of the gaze position (normalized with respect to the world-camera frame dimensions). Gaze data was recorded independently between the L and R eyes; for purposes of analysis we calculated the average gaze location between the two eyes for each timepoint.

### Analyses

#### Mapping to the target grid

For purposes of our analyses, it was necessary to transform all gaze data to be relative to the target grid stimulus. In practice we found that feature matching between the world-camera frames and the grid was improved if we embedded the grid within a larger, more visually complex border image. Thus, mapping the gaze positions to the target grid was a 2-step transformation: On each frame we first obtained and applied the transformation matrix mapping from the world-camera frame to the border image, and next applied the transformation mapping from the border image to the isolated calibration grid (the transformation matrix for this step needed to be determined only once, since the grid location was fixed with respect to the border image throughout the experiment. Moreover, given that this mapping occured between two hi-resolution, undistorted digital images, the transformation yielded negligible error). An alternative to this 2-step approach would be to simply hardcode the grid offset in pixels within the visually complex border image, and translate all of the mapped gaze locations accordingly.

#### Time-syncing gaze data and world-camera video frames

The eye-tracking data (i.e. world-camera video, gaze positions) and task data (i.e. trial onsets, trial order) were recorded by separate computers, and thus separate clocks. In order to synchronize the timestamps between the task and the eye-tracking data, the task included a brief visual cue prior to the first trial, presented by the task computer and presented in view of the eye-tracker world camera. Specifically, after calibration, the task computer was moved such that the screen was fully visible by the world-camera video stream. A red square appeared on the task computer for 3 seconds, and then was replaced by a visually complex start cue. Subsequent trial onsets were timestamped by the task computer relative to the presentation of this image. The feature-matching algorithm described above was used to find the frame in the world-camera video in which the start cue first appeared. By identifying the frame (and frame timestamp) in which this cue first appeared in the world-camera video, the gaze data could be aligned in time to the trial timestamps.

#### Measuring accuracy and precision

Each trial corresponded to one of the 9 total target points on which subjects were asked to fixate. All analyses took place on a window extending from 500-2500ms from trial onset. For each gaze position within that window, we calculated the distance (in degrees of visual angle) and polar angle relative to the target point (See **Fig 3.A**). Gaze points were excluded if they were more than 5° of visual angle away from the target in order to avoid outlier data points biasing accuracy measurements (“Accuracy and Precision Test Method for Remote Eye Trackers” 2011). We summarized the trial by calculating the mean gaze position across all raw gaze points on that trial (See **Fig 3.B**). Our **accuracy** measure was defined as the distance (in degrees of visual angle) between the mean gaze position and the target location (Holmqvist et al. 2011). Our **precision** measure was calculated as the square root of the arithmetic mean of squared distances between each raw gaze point and the mean gaze position (i.e. *standard deviation from mean gaze location,* or *root mean square (RMS) normalized by the mean).*

**Figure 3:**
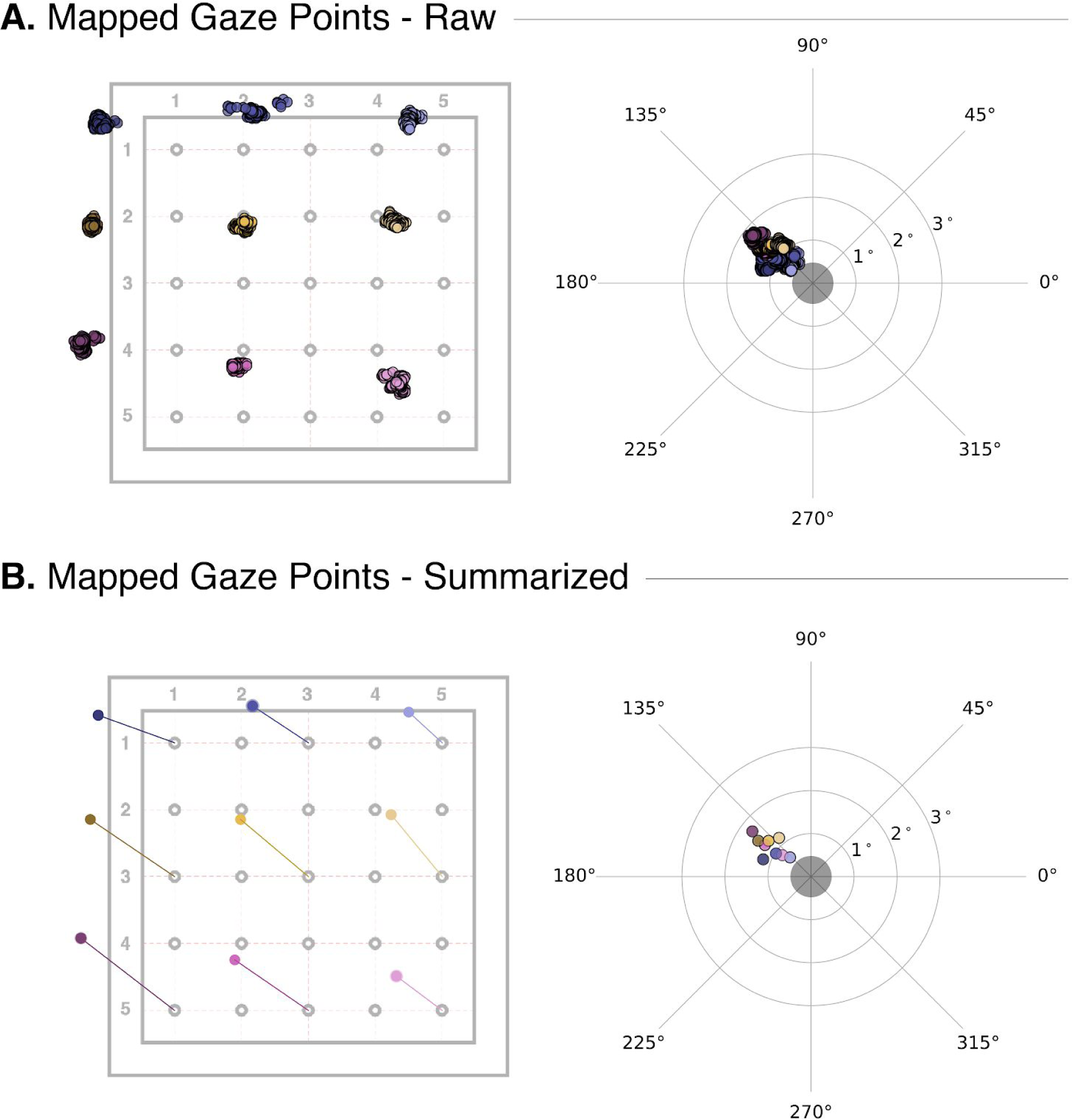
Sample data from a single task condition. Each condition consisted of 9 trials, during which participants were asked to fixate on a unique location within the target grid. The trial locations were presented in a random order, and for each location participants were asked to fixate on the point for 3000ms. **A.** Raw data was extracted from a window of 500-2500ms within each trial and mapped onto the target grid (left) as well as expressed in degrees of visual angle relative to each trial’s intended location (right). **B.**Visualization of the mean gaze location on each of the 9 trials. Distance and offset from the intended location is represented by lines (left), and degrees of visual angle (right).

*Summary plots for each subject and condition can be found at:*

*https://aithub.com/jeffmacinnes/alassesCalibration/tree/master/exposition/fias/conditionPlots*

## Results

*Code for all statistical models and result figures can be found online at: https://aithub.com/jeffmacinnes/alassesCalibration/blob/master/analvsis/calibrationAnalvses.md*

All analyses were completed using *R* v3.4.0 (R Core Team 2017), with data formatting using the *dplyr* package (Wickham et al. 2017). All linear mixed effects models were performed using the *lme4* package (Bates et al. 2015), and follow-up pairwise comparisons were performed using the *lsmeans* package (Lenth and Others 2016). All results plots were created using *ggplot2* (Wickham 2009), *ggpubr* (Kassambara 2017), and *ggsignif* (Ahlmann-Eltze 2017) packages for *R.*

### Overall Performance

The percentage of removed outlier (>5° from target location) gaze points during preprocessing differed by eye-tracker model. Statistical comparisons revealed the mean percentage of valid gaze points for Pupil Labs (mean: 97.1%; SE: 0.9) was significantly lower than SMI (mean: 98.7%; SE: 0.4) and Tobii (mean: 98.2%; SE: 0.4) (p < 0.001; no significant difference between Tobii and SMI). However, given that all models retained > 97% of all gaze points, this difference had a negligible effect on interpretation of subsequent analyses.

We first averaged across all distances and gaze angle conditions, and tested the overall relationships between eye-tracker model and accuracy, and eye-tracker model and precision.

### Accuracy

We fit a linear mixed effects model to test the relationship between accuracy and eye-tracker. This model included eye-tracker as a fixed effect and subject as a random effect. Eye-tracker was a significant predictor of accuracy (*F*(2,78) = 7.44, *p* < .001) in this model. Follow-up pairwise comparisons between eye-trackers revealed that the Pupil Labs eye-tracker was significantly more accurate than SMI (*t*(78) = 2.40, *p* < .05) and Tobii (*t* (78) = 3.81, *p* < .001). All other comparisons were non-significant at *p* > .1; see **Table 2**, and **Fig 4.A.**

**Figure 4:**
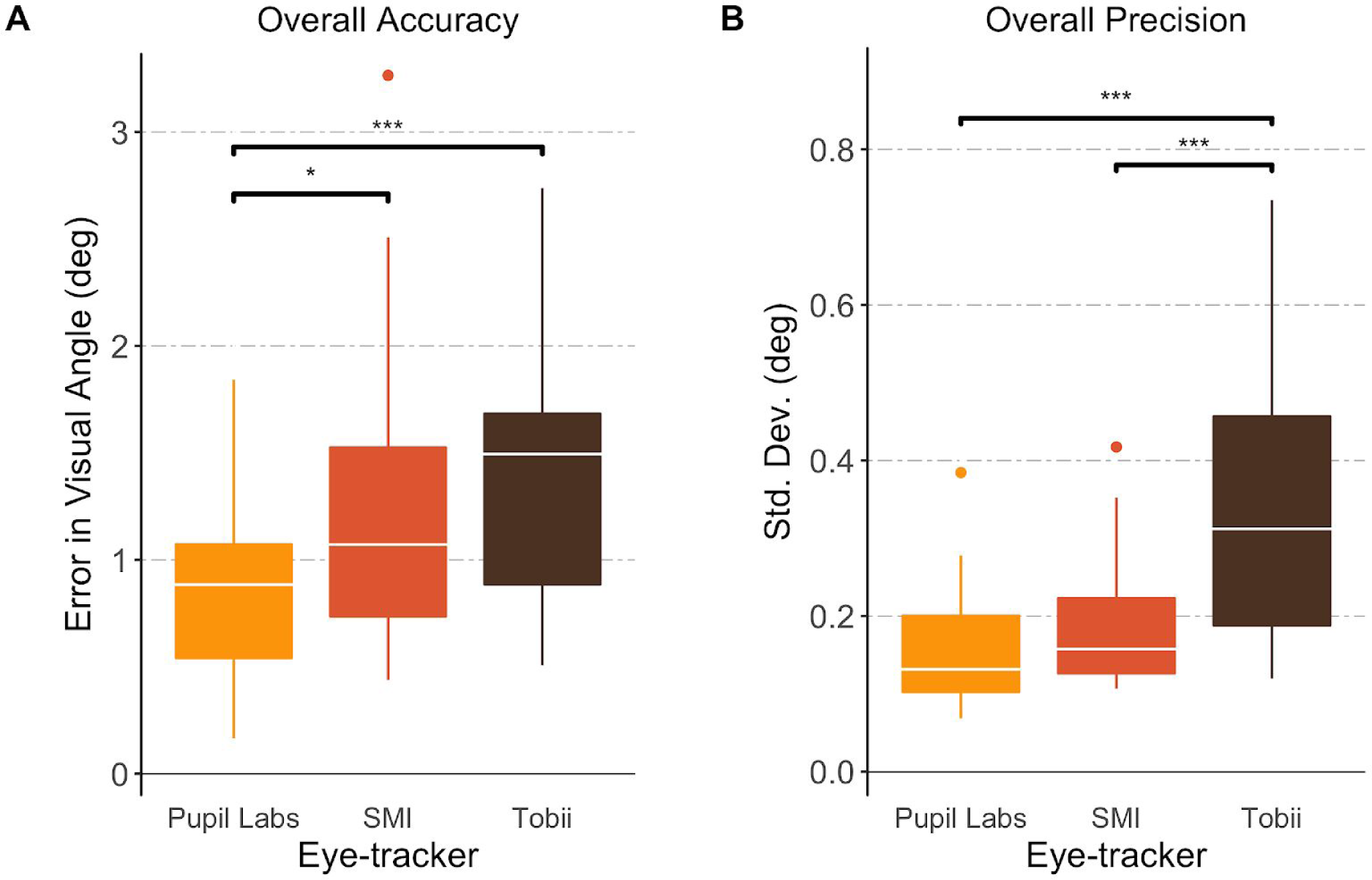
Overall Calibration Accuracy and Precision by Eye-tracker. **A.**Pupil Labs exhibited significantly less error (i.e. better accuracy) than either the SMI or Tobii eye-trackers, overall. **B.** The Tobii eye-tracker was significantly less precise, overall, than either the Pupil Labs or SMI eye-trackers.

**Table 1:** Wearable Eye-trackers. Device specifications across the 3 models of wearable eye-tracker tested in this study

| Model | World Cam.<br>Frame Rate | World Cam.<br>Frame size | World Cam.<br>FOV<br>(horizontal) | Gaze<br>Sampling<br>Rate | Calibration<br>Method |
| --- | --- | --- | --- | --- | --- |
| Pupil Labs 120<br>Hz Binocular | 30 *fps* | 1280 x 720 *px* | 60° | 120 *Hz* | ~9-pt Manual<br>marker calibration |
| SMI ETG 2.6 | 30 *fps* | 960 x 720 *px* | 60° | 60 *Hz* | 3-pt Natural<br>feature calibration |
| Tobii Pro<br>Glasses 2 | 25 *fps* | 1920 x 1080 *px* | 82° | 50 *Hz* | 1-pt Manual<br>marker |

**Table 2:** Overall accuracy and precision across wearable eye-tracker models

| Eye-Tracker Model | Mean Overall Accuracy | Mean Overall Precision |
| --- | --- | --- |
| Pupil Labs | 0.84° (0.42) | 0.16° (0.07) |
| SMI | 1.21° (0.66) | 0.19° (0.08) |
| Tobii | 1.42° (0.58) | 0.34° (0.16) |

### Precision

We fit a linear mixed effects model to test the relationship between precision and eye-tracker. This model included eye-tracker as a fixed effect and subject as a random effect. Eye-tracker was a significant predictor of precision (*F*(2,76) = 27.74, *p* < .001) in this model. Follow-up pairwise comparisons between eye-trackers revealed the Tobii eye-tracker was significantly less precise than Pupil Labs (*t* (76) = 6.96, *p* < .0001) and SMI (*t* (76) = 5.78, *p* < .0001). All other comparisons were non-significant at *p* > .1; see **Table 2**, and **Fig 4.B**.

### Differences by Distance and Offset

We next tested whether varying the distance and gaze angle influenced the accuracy and precision measures across the competing eye-tracker models.

#### Accuracy

We fit a linear mixed effects model to test the relationship between accuracy, eye-tracker, distance, and gaze angle. This model included eye-tracker, distance, and gaze angle as fixed effects and subject as a random effect. Our model examined main effects and interactions across all three of the fixed effects predictors. Fitting this model yielded a significant main effect of eye-tracker (*F*(2,54) = 8.69, *p* < 0.001), and a significant interaction between eye-tracker and distance (*F*(4,54) = 5.07, *p* < .01). Follow-up pairwise comparisons between eye-trackers at each distance revealed poorer accuracy on the Tobii eye-tracker compared to the Pupil Labs (*t*(54) = 5.23, *p* < .001) and SMI (*t* (54) = 2.99, *p* < .05) eye-trackers at a distance of 3*m* only. Pupil Labs compared to SMI accuracy was trend level (*t* (54) = 2.30, *p* = .06) at 3*m*. Lastly, there was also a trend level difference between Pupil Labs and SMI at a distance of *2m* (*t* (54) = 2.33, *p* = .06). All other comparisons were non-significant at *p* > .1; see **Fig 5**.

**Figure 5:**
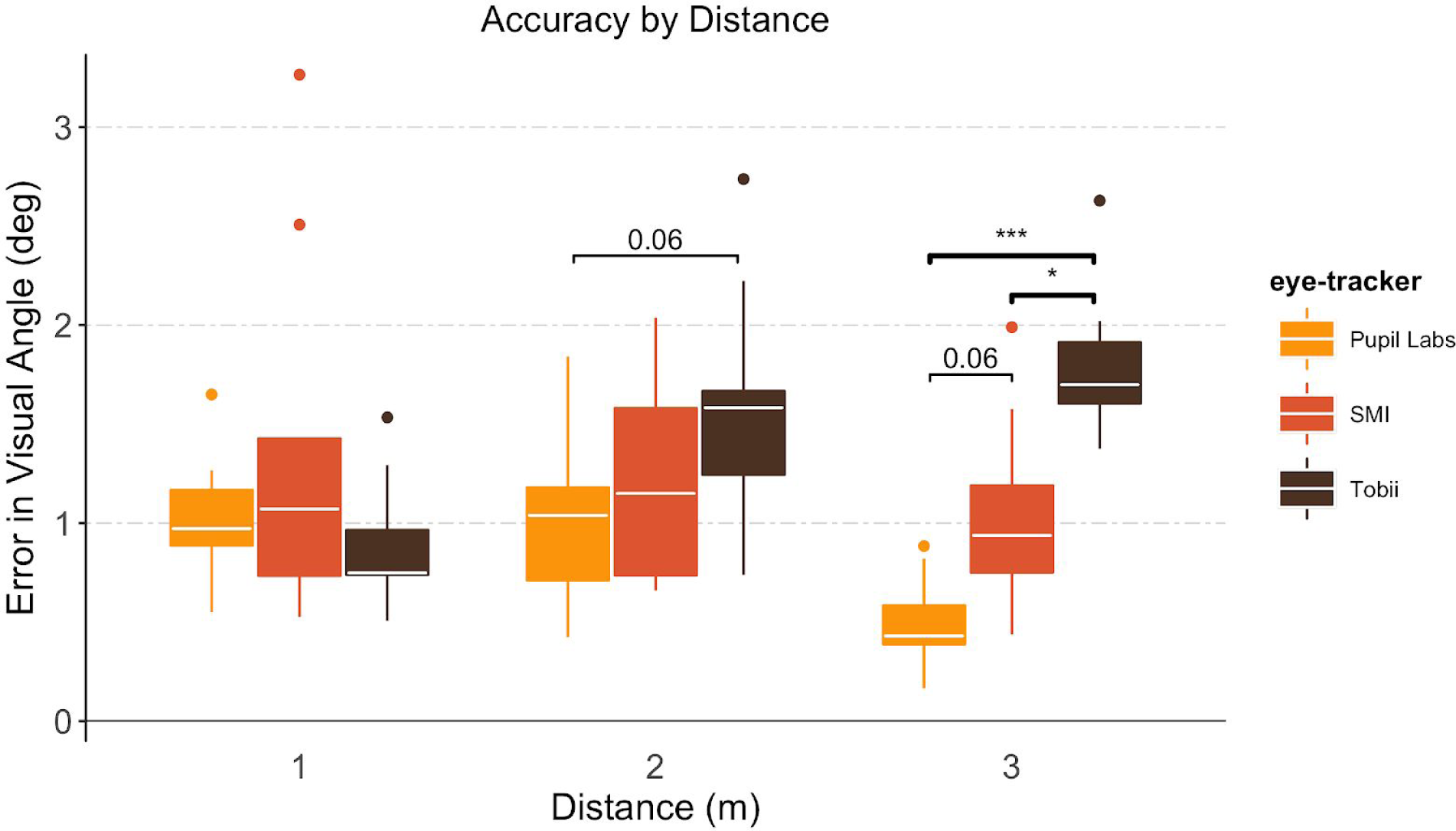
Influence of Viewing Distance on Calibration Accuracy by Eye-tracker. Differences in calibration accuracy were most pronounced at the farthest viewing distance (3*m*). At 3*m*, the Tobii glasses exhibited significantly worse accuracy than either the Pupil Labs or SMI eye-trackers. No significant differences in accuracy were observed at the shortest viewing distance (1*m*).

#### Precision

We fit a linear mixed effects model to test the relationship between precision, eye-tracker, distance, and gaze angle. This model included eye-tracker, distance, and gaze angle as fixed effects and subject as a random effect. Our model examined main effects and interactions across all three of the fixed effects predictors. Fitting this model yielded a significant main effects of eye-tracker (*F*(2,52) = 37.6, *p* < .001) and gaze angle (*F*(2,52) = 6.99, *p* < .01), and a significant interaction between eye-tracker and gaze angle (*F*(4,52) = 6.71, *p* < .001). Follow-up pairwise comparisons between eye-trackers at each gaze angle revealed the Tobii eye-tracker to be significantly less precise than the Pupil Labs at gaze angles of -10° (*t* (52) = 3.33, *p* < .01) and +10° (*t* (52) = 8.32, *p* < .001), and a trend toward less precise at the 0° condition (*t* (52) = 2.39, *p* = 0.05). In addition, the Tobii eye-tracker was significantly less precise than SMI at gaze angles of 0° (*t* (52) = 2.50, *p* < 0.05) and +10° (*t* (52) = 7.37, *p* < .001). All other comparisons were non-significant at *p* > .1; see **Fig 6**.

**Figure 6:**
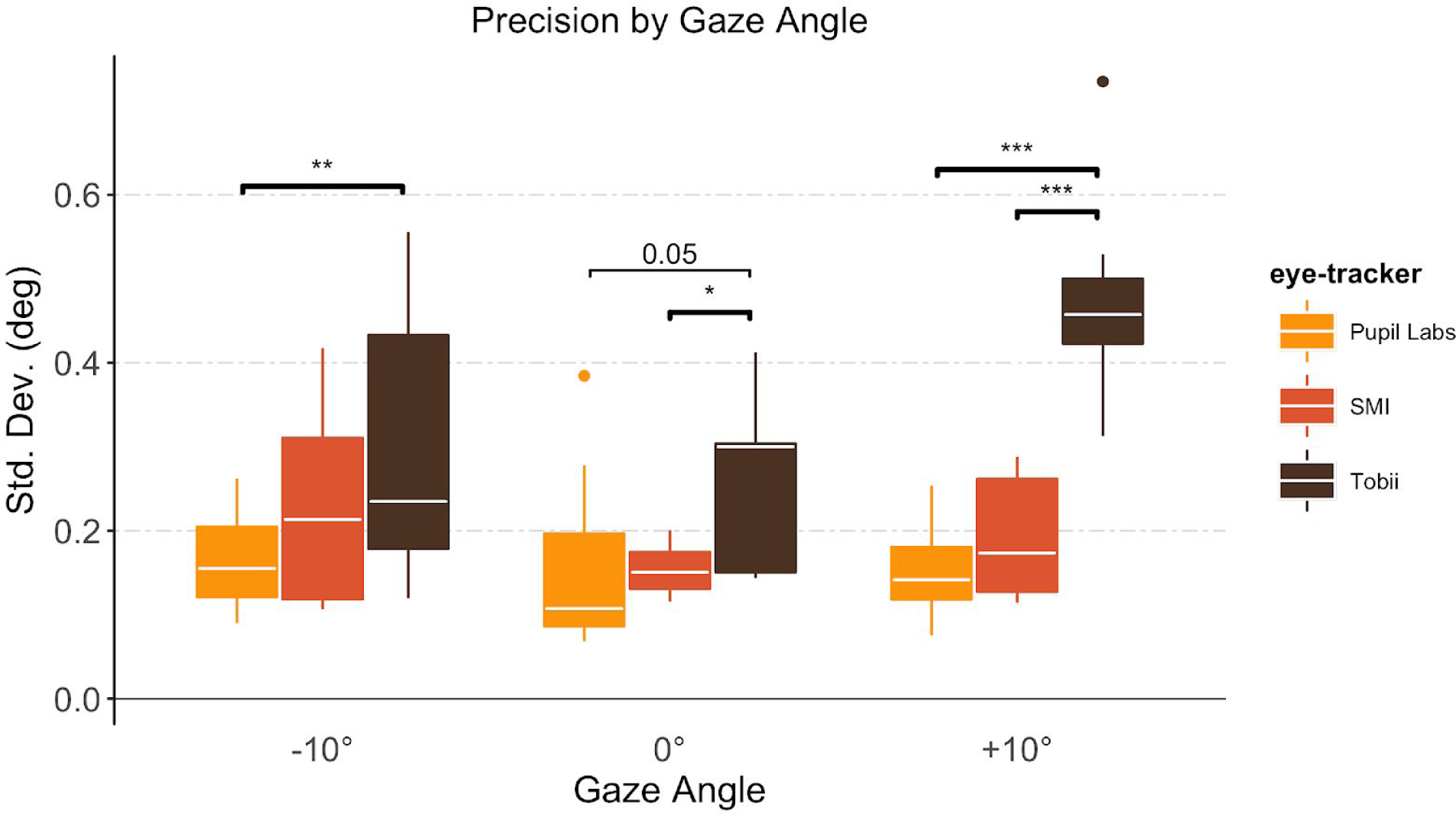
Influence of Gaze Angle on Calibration Precision by Eye-tracker. Pupil Labs exhibited better precision than the Tobii eye-tracker at gaze angles of +/-10°. A similar pattern was observed at 0°, albeit less robustly. Additionally, the SMI eye-tracker outperformed the Tobii eye-tracker at gaze angles of 0° and +10°. No significant differences in precision were observed between the Pupil Labs and SMI eye-trackers at any gaze angle.

## Discussion

### Dynamic Gaze Mapping

The freedom that wearable eye-trackers offer is achieved, in part, by reversing the frame of reference against which gaze positions are recorded. Whereas screen-based eye-tracking records gaze position with respect to the stimulus, wearable eye-trackers record gaze position with respect to the viewer. In other words, as a participant moves around, gaze positions are recorded on a fixed coordinate system that moves along with her/him. This introduces significant challenges to researchers attempting to study how participants view a particular stimulus over time.

In this article, we have detailed an automated approach to overcome the challenges that are introduced with wearable eye-trackers. Our pipeline uses freely available, open source software, built on top of feature matching algorithms developed for computer vision applications. Using this approach, researchers can dynamically map gaze position from a moving frame (i.e. the participant’s point of view) to a fixed reference coordinate system (i.e. the target stimulus). This approach makes it possible to study aggregate viewing behavior over a fixed 2D object in the environment, even as a participant continuously assumes new vantages points. We have made our specific software pipeline available for download, and note that this approach is easily adaptable to a wide variety of eye-tracking scenarios.

Nevertheless, there are caveats and limitations to this approach that should be considered. For one, this approach is limited in the types of target stimuli to which it can be applied. Target stimuli must be 2D (or approximated in 2D, e.g. a painting, a poster, the front surface of a cereal box), as the feature matching algorithm constructs a linear transformation that maps between two planar surfaces. Mapping gaze onto a 3D target (e.g. a sculpture) is a more computationally demanding challenge, but could be achieved by combining 3D reconstruction algorithms (Moons, Van Gool, and Vergauwen 2010; Lepetit and Fua 2005) with methods for estimating participants’ gaze depth (Mansouryar et al. 2016; Lee et al. 2012). Furthermore, ideal targets should contain non-symmetric, unique elements with relatively high-spatial frequency information, in order to ensure a sufficient set of keypoints can be identified on the reference image. This approach would likely fail, for instance, with minimalist paintings such as the large monochrome works of Yves Klein. Similarly, our approach is best suited for 2D targets that are static and visually varied. In principle, the same approach could be used for 2D targets that change over time (e.g. a video clip); however, unlike the current implementation, in which we obtain keypoints on the reference target image once at the beginning, video would require obtaining a unique set of reference keypoints for each frame of the target video (and both the target video and world camera video would have to be perfectly synchronized).

We observed no appreciable differences in how well the SIFT feature-matching algorithm performed across the 3 eye-trackers we tested. The factor most likely to influence this would be the degree of lens distortion on the world camera. The gaze mapping algorithm relies on a linear transformation between the world camera video frame and the reference image; camera lenses, necessarily, introduce non-linear distortions to the frame that adversely affect the precision of that transformation. Distortions are most evident along the periphery of the frame and are often more severe with wider-angle lenses. As long as distortions are minimized (or corrected via preprocessing steps), our approach would be expected to perform similarly well with other eye-trackers. We present validation of this approach under conditions where the target stimulus is between 1-3m distant and within +/-10° of viewing angle. This is a subset of the full range of viewing conditions that researchers may wish to employ, but one that should encapsulate a wide range of indoor viewing contexts. Moreover, while our validation data presented results from participants in a fixed position, our gaze mapping approach is robust to changes in head motion; since gaze mapping proceeds independently on each frame from the world camera, it is unaffected by frame-to-frame changes in position. The caveat to this is that slow frame rates on the world camera would be less susceptible to motion blur, which can adversely affect the ability to detect matching key points due to blurred stimuli. Conversely, faster frame rates help mitigate this issue.

The approach we have described is robust, and can map gaze position automatically with no manual oversight required. Nonetheless, there are numerous ways to optimize the computations and thereby shorten the processing time (anecdotally, on a machine with a 2.6 GHz processor and 16Gb of RAM, our approach takes ~1.5s/frame, or ~90 min for a 2 min recording at 30 *fps*). Our algorithm runs serially, but independently, on each frame of the world camera recording. One way to improve the processing time would be to simply parallelize the algorithm to execute multiple frames simultaneously. We also note that treating the frames independently fails to take advantage of sequential aspects of the recording. Methods for efficient pixel tracking in video, like optical flow (Lucas, Kanade, and Others 1981), can be utilized to substantially reduce the search space for keypoints on consecutive frames (Ta et al. 2009), greatly reducing processing time. We leave these optimization steps to future work.

### Comparing Wearable Eye-trackers

Researchers interested in taking advantage of the benefits of wearable eye-trackers face the difficulty of choosing between competing devices on the market. Existing models vary in specifications, features, and price. While choosing the most appropriate eye-tracker involves many factors and depends on the unique needs of a research group, we designed a task to benchmark basic accuracy and precision performance under conditions that approximate how these devices might be used in indoor, short-to-medium viewing range, research settings. Using this task, we compared performance across 3 different available models of wearable eye-tracker: Pupil Labs 120 Hz Binocular, SMI ETG 2.6, and Tobii Glasses 2.

Overall, the Pupil Labs 120Hz Binocular glasses exhibited significantly smaller accuracy errors (average error of < 1°) than the SMI and Tobii models. The Pupil Labs eye-tracker also exhibited significantly tighter overall precision than the Tobii eye-tracker (no significant difference in overall precision between the Pupil Labs and SMI eye-trackers).

We analyzed these results further by asking whether eye-tracker performance would vary as a function of distance or gaze angle. Our task spanned a parameter space of 3 distances (1*m*, 2*m*, 3*m*) and 3 gaze angles (-10°, 0°, +10°) in order to test performance at the type of short- to medium-range viewing conditions that may be applicable to researchers. We found that accuracy differences between eye-trackers was influenced by the distance at which they were tested. Namely, the biggest differences in accuracy emerged at the farthest testing distance (3*m*). At that distance, the Tobii eye-tracker was significantly less accurate than the Pupil Labs eye-tracker, and marginally less accurate than the SMI eye-tracker. Notably, there were no differences in accuracy among the eye-trackers at the closest distance (1*m*).

In terms of precision, distance was not a significant predictor, but gaze angle was. We found that the Pupil Labs eye-tracker exhibited tighter precision than the Tobii eye-tracker across all three gaze angles (note: this difference was only marginally significant at 0°), and that the SMI eye-tracker outperformed the Tobii eye-tracker at gaze angles of 0° and +10°. The Pupil Labs and SMI eye-trackers exhibited similar precision performance across all gaze angles. It is not clear from the design of the eye-trackers why precision would be different at -10° compared to +10°. While the differences between Tobii and the other two eye-trackers were most pronounced at +10°, we note our comparisons did not directly test for differences within the same eye-tracker across gaze angles. In general, the patterns of differences we observed at each gaze angle are consistent with the overall differences in precision between eye-trackers.

It is worth highlighting that the accuracy and precision measures we analyzed in this study depend critically on the initial calibration steps that each eye-tracker performed. In other words, the accuracy of our automated gaze mapping approach will only be as good as the manufacturer-specific calibration between the recorded gaze location and the participant’s actual gaze location. The calibration procedures vary across the eye-trackers we tested, as do the number of parameters the experimenter can control. We note that accuracy and precision performance in this study was inversely correlated with the how adaptable the calibration procedures were to the particular viewing conditions. The Pupil Labs eye-tracker recommended calibrating using ~9 separate calibration points, positioned at the same distance as data collection for each condition. The 9-pts allowed us to verify accurate calibration across a wide portion of the participant’s field of view. By contrast, the Tobii recommended procedures were to use a single point held at a distance of ~1*m* away. Calibration procedures will only yield accurate data at the distance at which calibration occurs. Indeed, we observed no significant differences in accuracy between any of the eye-trackers at 1*m*. We suspect that each manufacturer weighed the tradeoff between user friendliness and optimization when deciding which calibration options to make available. While Pupil Labs offered the most optimization options, it also took the most time to set up and calibrate. On the other hand, while the Tobii model was the easiest to set up, our experience suggests that the accuracy performance (particularly at farther viewing distances) could be improved with more flexible calibration procedures.

Thus, for close-range viewing experiments at 1*m* or less (e.g. participants interacting with handheld tablets), there may be no appreciable performance differences across devices. However, as viewing distance increased, performance differences between the devices emerged. Our findings suggest that enhancing the initial calibration procedures would likely yield more consistent results across devices. We also note that other variables, such as cost and ease-of-use, may be important factors in choosing the appropriate eye-tracker.

### Implementing Wearable Eye-trackers in a Research Context

As noted above, obtaining a good initial calibration is crucial to ensuring accurate results during subsequent steps of the analysis. Although each device we tested used its own unique calibration procedures, there are generalized steps that apply across devices, and will likely apply to future models that are released. First, we found that the best way to ensure a good calibration is to verify it immediately afterwards. This typically involved asking the participant to fixate on specific locations (e.g. the calibration target itself) and visually verifying that the projected gaze location in the live view within eye-tracker software matched reasonably well. We typically asked participants to keep their head still, and then fixate at 3-5 distinct locations that spanned the field of view. If the projected gaze locations failed to align with the actual location, we repeated the calibration procedure until this improved. With the Pupil Labs eye-tracker, this process could take as many as 8-10 iterations, perhaps because of the comparatively large number of calibration points (9).

In addition, we also found that performance increased with frequent re-calibrations. In our study we re-calibrated participant’s before each unique condition, which meant each calibration was good for ~1-2 mins of data recording. Each of the devices we tested purports to build a 3D model of the eyeballs during calibration. This model allows the devices to compensate for movement or slippage of the glasses that occurs during normal use so as to preserve the calibration mapping. While these models surely helped, we nevertheless found that the projected gaze location would drift from the intended target over time. In most cases, this was corrected by simply re-calibrating the eye-tracker regularly.

Lastly, the accuracy of the gaze mapping is only as good as the linear transformation between the world camera video and reference image on each frame. Obviously, gaze data cannot be mapped to a reference image when the participant is not oriented toward the target stimulus (impossible to use feature-matching if there are no matches on the world camera video frame).

Less obviously, gaze mapping performance deteriorates as the frame gets blurrier (during periods of fast motion, for instance). This effect can be mitigated by faster frame rates with the world camera, or instructing participants to move more slowly.

## Conclusion

Screen-based eye-tracking methods impose considerable constraints on both participants and stimuli, presenting major impediments for naturalistic viewing studies. By contrast, wearable eye-trackers, which have recently become available, allow researchers to record gaze behavior as participants freely navigate an environment and interact with real-world stimuli. Despite these advantages, adapting these devices to research questions presents numerous challenges. Analyzing data from wearable devices is complicated by the fact that gaze position is recorded in reference to a participant’s point of view, rather than in reference to the target stimulus in the environment. In this article we outlined an approach to automatically and dynamically map gaze data to a fixed reference image, facilitating the use of this methodology in experimental contexts.

Pursuing this methodology can be a difficult decision for interested researchers, in part because of the significant costs involved and the lack of comparative performance measures between competing devices on the market. We addressed this issue by developing a method to benchmark the accuracy and precision of wearable eye-trackers, and tested this method against existing models of wearable eye-trackers. Of the models we tested, the Pupil Labs 120Hz Binocular glasses exhibited the least accuracy error overall. The Pupil Labs 120Hz and the SMI ETG 2.6 displayed similar precision performance overall, with both showing tighter precision than the Tobii Pro Glasses 2. However, viewing distance and gaze angle were important factors in interpreting our results.

Wearable eye-trackers offer a substantial advantage to studying naturalistic viewing behavior; we hope the our approach and results will assist other research groups.

## Acknowledgments

The authors extend their gratitude to Kathryn Dickerson, Shad Todd, Marianne Wardle, Erik Wing, and Jonathan Winkle for their assistance in data collection, and to Robert Botto for insightful comments and editing. This work was supported by Duke University’s Bass Connections - Brain & Society, and the Big Data to Knowledge Initiative (K01-ES-025442) awarded to JP.

